# Modeling Thoracic Aortic Dissection Using Patient-Specific iPSCs Reveals VSMC Dysfunction and Extracellular Matrix Dysregulation

**DOI:** 10.1101/2024.10.06.616859

**Authors:** Peifeng Jin, Yubin Xu, Sixian Wang, Lu Ding, Yuhao Chen, Miqi Zhou, Xiufang Chen, Xiaofang Fang, Yongsheng Gong, Ming Li, Yongyu Wang

**Affiliations:** Department of Cardiac Surgery, the First Affiliated Hospital of Wenzhou Medical University, Wenzhou 325000, Zhejiang, China; Institute of Hypoxia Medicine, School of Basic Medical Sciences, Wenzhou Medical University, Wenzhou 325015, Zhejiang, China; Department of Biochemistry and Molecular Biology, School of Basic Medical Sciences, Wenzhou Medical University, Wenzhou 325015, Zhejiang, China; Cardiac Regeneration Research Institute, School of Basic Medical Sciences, Wenzhou Medical University, Wenzhou 325015, Zhejiang, China

**Author notes:** These authors contributed equally to this work and share first authorship (Peifeng Jin, Yubin Xu). Correspondence: Yongyu Wang.

**Keywords:** Thoracic Aortic dissection, iPSCs, VSMCs, Disease modeling, collagen IV

## Abstract

Thoracic aortic dissection (TAD) is a life-threatening condition characterized by medial degeneration and vascular smooth muscle cell (VSMC) dysfunction, with no effective medical therapy currently available. The underlying pathological mechanisms of TAD remain incompletely understood. In this study, we used a non-integrated episomal vector-based reprogramming system to generate induced pluripotent stem cells (iPSCs) from TAD patients and healthy controls. Both TAD and normal iPSCs expressed key pluripotency markers and were capable of differentiating into the three germ layers in vitro. These iPSCs were differentiated into VSMCs through a mesodermal intermediate for disease modeling. VSMCs derived from both TAD and normal iPSCs expressed smooth muscle α-actin (α-SMA), calponin, and SM22α. However, TAD-iPSC-derived VSMCs exhibited significantly reduced contraction in response to carbachol stimulation compared to their normal counterparts. Whole-exome sequencing identified a mutation in the COL4A2 gene (c.392G>T, p. R131M) in TAD-iPSCs. This mutation was associated with reduced collagen IV expression and increased expression of collagen I and III in TAD-VSMCs, both with and without TGF-β stimulation. Furthermore, noncanonical TGF-β signaling was hyperactivated in TAD-VSMCs, accompanied by elevated MMP9 expression. This patient-specific iPSC model reveals key dysfunctions in VSMC contractility, extracellular matrix protein expression, and dysregulated TGF-β signaling, which may contribute to TAD pathogenesis. Our findings provide new insights into the molecular mechanisms driving TAD and offer a platform for future therapeutic development.

## 1 Introduction

Thoracic Aortic dissection (TAD) is an aggressive and life-threatening vascular disease with high mortality^1–3^, affecting approximately 3-4 cases per 100,000 annually^4^. This condition is characterized by a tear in the aortic wall, leading to the separation of the layers of the aorta, which can result in severe complications such as rupture or organ damage. The pathogenesis of TAD is complex and multifactorial. Major risk factors include hypertension, dyslipidemia, aging, and genetic predisposition. Genetic analysis has revealed that over 20% of individuals with TAD or thoracic aortic aneurysm (TAA) – defined as a dilation of the thoracic aorta to a diameter at least 1.5 times greater than normal at a given aortic level – possess single-gene mutations^5^. To date more than 30 genes associated with TAD and TAA have been identified, emphasizing the genetic component of this disease ^6^. Current understanding of TAD pathogenesis is limited, and no effective medical therapies are available. The standard treatment involves surgical intervention, which carries significant risks and limitations^7^. The lack of effective pharmacological treatments underscores the urgent need for novel therapeutic strategies and a deeper understanding of the disease mechanisms. Although extracellular matrix degradation and inflammation have been suggested as the key features of TAD, the precise mechanisms still need to be characterized^2, 8^.

The aorta wall is composed of three layers the tunica intima, tunica media, and tunica adventitia. The media of the aortic wall consists of smooth muscle cells and extracellular matrix (ECM), which includes elastic fibers, collagen, proteoglycans, glycosaminoglycans, and various adhesive proteins. Maintaining ECM homeostasis is a critical for normal aortic function, and dysregulation of ECM is associated to TAD, characterized by medial degeneration. Collagen is a crucial component of the ECM, essential for maintaining the structure and function of the aorta. Type I and III collagen are the most abundant collagen present in the aortic wall and show increased expression and deposited in TAD. Mutations in the *COL3A1* gene are linked to vascular-type Ehlers-Danlos syndrome, leading to abnormal collagen fibrillogenesis and TAD formation^9^. Additionally, mice with *col1a1* mutations also exhibit age-dependent onset of aortic dissection^10^. Conversely, the type IV collagen, a basement membrane-specific collagen, is expected to have decreased expression in TAD^11^. Although mutations in collagen-encoding genes are closely associated with aortic dissection, the underlying mechanisms remain unclear.

VSMCs play an essential role in the pathogenesis of aortic dissections^12^. The primary function of VSMCs in blood vessels is to regulate blood pressure and blood flow distribution. VSMCs can undergo phenotypic switching in response to various pathological conditions, such as atherosclerosis, hypertension, restenosis. In these cases, contractile SMCs switch to synthetic phenotype, characterized by reduced the expression of contractile and cytoskeletal proteins, increased proliferation and migration, and overproduction of components involved in aortic wall remodeling, such as matrix metalloproteinases (MMPs) ^13^. The loss of the contractile phenotype of VSMCs is recognized as an early event in thoracic aortic aneurysm^14^. However, whether the phenotypic switching of VSMCs mediates collagen mutation-induced aortic dissection remains unclear.

Recent advances in stem cell technology offer promising avenues for disease modeling and therapeutic development. Induced pluripotent stem cells (iPSCs), which are generated by reprogramming somatic cells, have the potential to differentiate into various cell types and provide valuable insights into disease mechanisms at the cellular level. iPSCs can be derived from patients with specific genetic conditions, allowing the creation of patient-specific disease models^15^. These models can be used to study the molecular and cellular bases of various diseases including cardiovascular disease and in testing potential therapeutic agents^16, 17^. For instance, Granata et al. successfully used Marfan syndrome-iPSC to model the key mechanism of smooth muscle cell death in Marfan syndrome^18^. Similarly, generation of patient-specific iPSC from TAD patients in this study may serve as a new powerful tool to study the molecular mechanisms of TAD *in vitro*.

In this study, we employed a non-integrated episomal vector-based reprogramming system to generate iPSCs from TAD patients and healthy controls. Our aim was to investigate the cellular and molecular mechanisms underlying TAD using these patient-specific iPSCs. By comparing the properties and behaviors of TAD patient-derived iPSCs with those from healthy individuals, we observed that TAD-iPSC-derived VSMCs exhibited dysfunction, including reduced contractile response and elevated cell proliferation rates. Additionally, a COL4A2 mutation was identified in TAD-iPSC, which showed decreased expression of type IV collagen and increased expression of type I and III collagen. Furthermore, we found that the ERK-mediated noncanonical TGF-β pathway was activated while the canonical TGF-β-Smad pathway was repressed, indicating that TGF-β is a key pathway and a potential therapeutic target involved in TAD pathogenesis.

## 2 Materials and methods

### 2.1 Isolation and culture of primary aortic SMCs

The aortic tissues from TAD were obtained from the Department of Cardiothoracic Surgery at the First Affiliated Hospital of Wenzhou Medical University. The aortic specimens were washed twice with HBSS (Invitrogen). The endothelial layer and the tunica adventitia were removed, and tunica media was cut into pieces of about 1 mm3 explants and were placed on a flask for overnight culture with a small amount of Dulbecco’s Modified Eagle’s Medium (DMEM) with 10% fetal bovine serum and 1% penicillin/Streptomycin at 37 °C with 5% CO2 as a previous report^19^. The medium was replenished to cover the explants on the next day. The medium was changed every 3-4 days till the outgrown cell monolayer reached 90% of confluence (normally taking 2-3 weeks). The primary cells were then passaged for iPSC reprogramming.

### 2.2 Generation of human iPSCs

The non-integrative iPSCs were generated by nucleofecting TAD-VSMC or BJ cells (normal skin fibroblasts) with episomal vectors (total 3 μg; pCXLE-hUL, pCXLE-hSK, pCXLE-hOCT3/4-shp53-F, and pCXWB-EBNA1) by AMAXA 4D Nucleofector (LONZA) as previously described ^20^. Six days post-nucleofection, 2×105 cells were replated on a matrigel-coated 35 mm dish with DMEM containing 10% FBS. After 24 h of replating, the medium was switched to the human reprogramming medium (ReproEasy, Cellapy, CA5001500) and changed every other day. Two weeks after transduction, the medium was changed daily with hESC medium until iPSC colonies were observed. The generated iPSCs colonies were picked manually and maintained in feed-free matrigel-coated dishes, The iPSCs were fed every other day and were passaged 1:5-10 every 5 days using Versene (Invitrogen). All the protocol was approved by the Medical Research Ethics Committee of Wenzhou Medical University (2017-066) and written informed consent was obtained from the patients.

### 2.3 Characterization of iPSCs (undifferentiation and multi-differentiation)

Alkaline Phosphatase (AP) Staining Kit II (Beyotime, C3206) was used for AP staining of iPSCs as the manufacturer’s instruction. Briefly, iPSCs were fixed with 4% paraformaldehyde at room temperature for 5 min after PBST wash. Then cells were incubated in AP Substrate Solution for 15 minutes, to stop the reaction by washing with PBS. For immunofluorescence staining, iPSCs or differentiated cells were fixed in 4% paraformaldehyde for 20 minutes, permeabilized and blocked using 0.5% Triton X-100 in PBS with 5% normal goat serum, and incubated overnight with the primary antibodies: rabbit anti-OCT4, rabbit anti-SOX2, rabbit anti-Nanog, mouse anti-TRA-1-60, mouse anti-TRA-1-81 (1:100, Stemgent), mouse anti-TUJ1 (1:500, Covance), rabbit anti-α-1-fetoprotein (1:400, Abcam), mouse anti-αSMA (Sigma), mouse anti-calponin (Sigma), rabbit anti-smooth muscle 22α (SM22α) (Abcam). Secondary goat-anti-mouse and anti-rabbit IgG antibodies conjugated with Alexa 488 or Alexa 594 were added to the samples and incubated for 1 hour. The cell nuclei were stained with 4′,6-diamidino-2-phenylindole (DAPI) (Life Technologies). Cells were rinsed, and fluorescence was analyzed using a Nikon (Ts2-FL) microscope.

### 2.4 Teratoma formation assay

Immunodeficient SCID mice were injected 1×106 iPSCs with DMEM/Matrigel solution into the hind limbs intramuscularly. Eight weeks after injection the teratomas were dissected and fixed with 4% paraformaldehyde. Paraffin-embedded tissue was sliced and stained with hematoxylin and eosin.

### 2.5 SMC differentiation from iPSCs

SMC differentiation from iPSCs was performed according to our previous human ES cells protocol with minor modification^21^. iPSCs were digested with 1 mg/ml collagenase IV (Solarbio) for 15-20 min and transferred to the ultra-low-attachment dish (corning) to make embryoid bodies (EBs). EBs were cultured in a human ES cell medium without bFGF. Six-day-old EBs were replated to 35mm dishes coated with 0.1% gelatin and cultured for 6-12 days in DMEM with 10% FBS. The outgrowth cells from EBs were dissociated with TrypLE express and cultured on Matrigel-coated 35mm dishes with smooth muscle growth medium2 (SmGM2) (Lonza). Cells were passaged when they reached 80%–90% confluence. To induce differentiation to SMCs, cells were cultured on differentiation conditions, in which cells were grown on gelatin-coated dishes in SmGM2 basal medium with 5% FBS for 5 days.

### 2.6 Contractility assay

After 5 days of differentiation, SMCs were washed with phosphate-buffered saline (PBS), stimulated with 1 mM carbachol in smooth muscle differentiation medium, and monitored for 30 minutes as we described before ^21^. Images of the same field were collected every 1 minute for 30 minutes and compiled into movies using the Olympus software (DP controller). To analyze the change in cell surface area between 0 min and 30 min using image J software.

### 2.7 Quantitative RT-PCR

Total RNA was extracted using the RNeasy Mini kit (Qiagen). cDNA was synthesized from 1 µg of RNA with a PrimeScript RT reagent Kit (Takara, RR037A). Pluripotency genes were amplified with gene-specific primers by PCR using Taq DNA polymerase (NEB, M0267V) in SimpliAmp™ Thermal Cycler (Applied Biosystems). Real-time PCR was performed on a StepOne Plus Real-Time System (Life Technologies) using iTaq Universal SYBR Green Supermix (Bio-Rad) as previously described^22^. Quantification of gene expression was assessed with the comparative cycle threshold (Ct) method. The relative amounts of mRNA for the different genes were determined by subtracting the Ct values for these genes from the Ct value for the housekeeping gene GAPDH (ΔCt). Primer sequences are shown in supplemental online Table 1.

### 2.8 Flow cytometry

Cells were dissociated with 0.05% trypsin into single cells and suspended in PBS, then fixed with the fixation solution from the Fixation/Permeabilization kit (BD Biosciences), and stained with primary and detection antibodies as described by the manufacturer. Briefly, the dissociated cells were resuspended (5 ×105 cells) in 250 μL of fixation/permeabilization solution, kept at room temperature for 30 min, and washed twice with Perm/Wash buffer. After blocking with blocking solution for 30 min, the cells were incubated with the primary antibody for 1h on ice. The cells were then resuspended in 100 μl of Perm/Wash buffer after incubation with a secondary antibody for 1h on ice, washed twice, and analyzed by flow cytometry. The primary and detection antibodies used are listed in Table 2.

### 2.9 Karyotyping

Karyotyping of iPSC was performed on G-band metaphase chromosomes by Nuwacell Ltd. The proliferating iPSCs were blocked by 50 ng/ml of colcemid for 2 h, digested with trypsin-EDTA, and then treated with hypotonic KCl solution for 20-40 min at 37 °C. Glass slides were prepared with three steps of fixation in methanol/glacial acetic acid (3:1). QFQ-banding was analyzed at a resolution of 400 bands per haploid genome according to the International System for Human Cytogenetic Nomenclature (ISCN2016).

### 2.10 Western blotting analysis

Cells were lysed in RIPA buffer on ice and protein concentrations were determined by BCA protein assay (Thermo Scientific, Rockford, IL) according to the manufacturer’s protocol. 30 ug of cell lysate was resolved by 10% sodium dodecyl sulfate (SDS)-polyacrylamide gel and transferred to nitrocellulose (NC) Protran membranes (Whatman, Dassel, Germany). Blots were incubated for 1 hour at RT in blocking buffer (5% milk in tris-buffered saline (150 mM NaCl, 10 mM Tris, pH8.0, 0.1% Tween-20)), and with primary antibody overnight at 4°C, followed by horseradish peroxidase-conjugated secondary antibody (1:20,000 in blocking buffer) for 1 hour at RT, and visualized by SuperSignal West Pico Chemiluminescent substrate (Thermo Scientific, Rockford, IL) and were detected using a digital gel image analysis system (Bio-Rad, Hercules, CA, USA). The density of the protein bands was normalized to the GAPDH and presented as a percentage increase.

### 2.11 Statistics

Statistical analyses were performed using Prism (GraphPad Software, Inc., La Jolla, CA). Numerical data were reported as mean ± s.e.m. From triplicate cell culture (n = 3), the Student’s t-test was applied to test the significance between the groups. The value of p < 0.05 was considered statistically significant.

## 3 Results

### 3.1 Generation and characterization of non-integrated iPSCs from TAD patient

The generation of integration-free iPS cells is crucial for disease modeling, cell-based therapy, and regenerative medicine. Using our previously established the episomal vector-mediated human iPSCs reprogramming platform, we generated non-integrative iPSCs from a 47-year-old patient with thoracic aortic dissection (TAD) (Fig.1 and Fig.S1). Additionally, we also generated a normal iPSC from BJ cells, which are normal human dermal fibroblasts (Fig.1 and Fig.S1). We picked different colonies to establish at least three lines for each sample based on their ESC-like morphology. We characterized the pluripotency of these iPSC lines using alkaline phosphatase staining (APS) and immunofluorescence. Fig.1B showing a representative TAD-iPSC and normal iPSC lines with positive APS staining and expression of pluripotency markers, including the transcription factor OCT4, NANOG, SOX2, as well as the cell surface protein TRA-1-60, TRA-1-81. RT-PCR analysis confirmed that the pluripotency genes of OCT4, NANOG, SOX2 were highly expressed in all the iPSC lines (Fig.1C). Karyotyping assays confirmed that both the normal iPSCs and TAD-iPSCs exhibited a normal karyotype (Fig.1D). To further characterize the pluripotency of normal and TAD-iPSCs, we assessed their differentiation potential into the three germ layers in vitro. We used antibodies for α-fetoprotein (AFP), smooth muscle α-actin (α-SMA), and TUJ1 to determine endoderm, mesoderm, and ectoderm differentiation, respectively. As shown in Fig. 1E, both normal and TAD-iPSC lines could differentiate into three layers in vitro. When these iPSCs were inoculated into SCID mice, they formed teratomas in vivo. Histological staining of these teratomas indicated the presence of various tissues representing all germ layers, including neural rosette (ectoderm), muscle (mesoderm), and gut-like epithelial (endoderm) (Fig.1F).

**Fig. 1.**
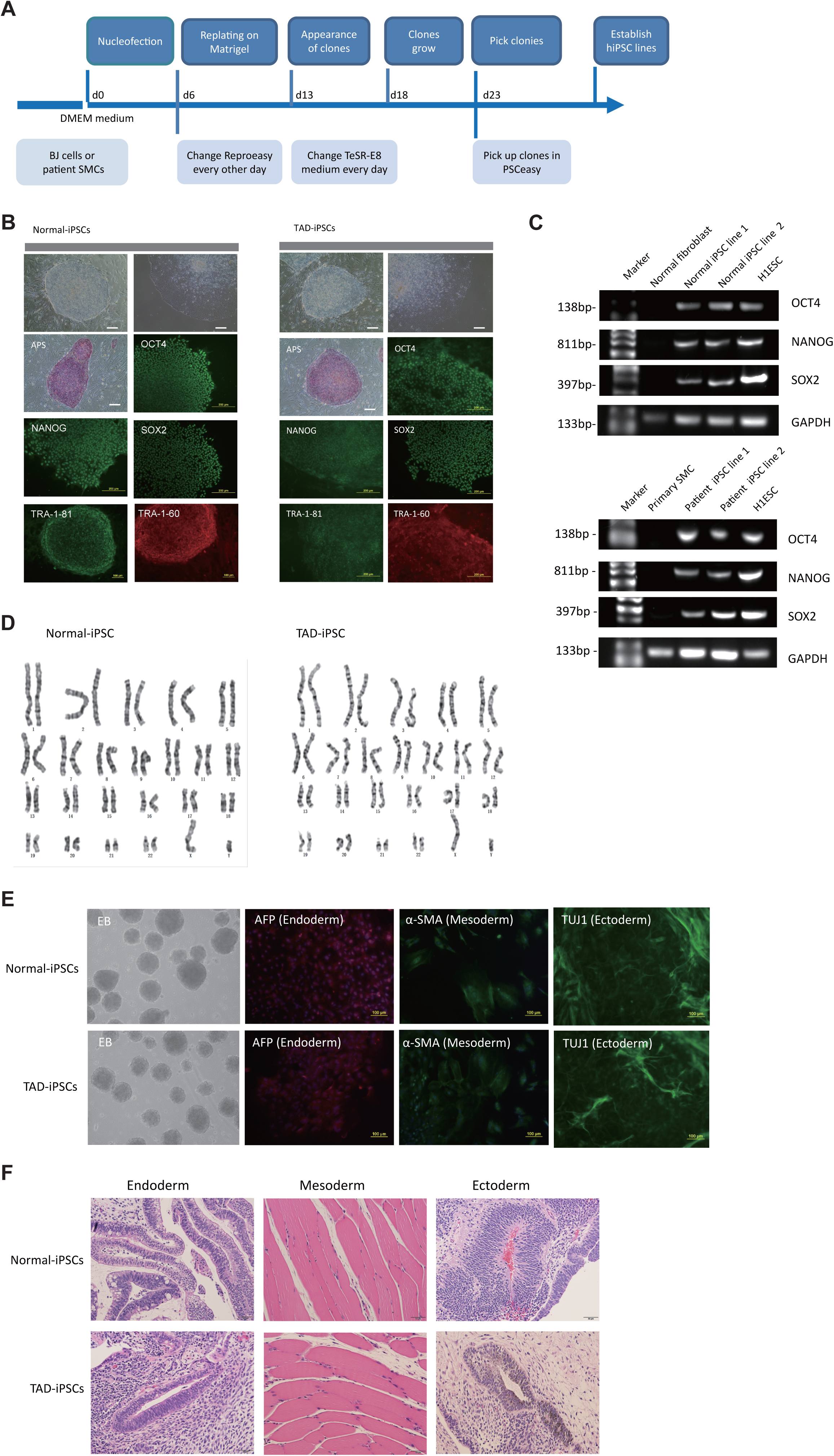
Generation and characterization of hiPSCs from TAD patients and normal cells. (A) Schematic diagram of non-integrative iPSCs reprogramming mediated by episomal vectors. (B) The morphology of established iPSC clone, alkaline phosphatase staining (APS), and immunofluorescence staining with pluripotency markers, including, OCT4, NANOG, SOX2, TRA-1-60, TRA-1-81. Scale bar=200μm. (C) Pluripotency marker gene expression assessed by RT-PCR. (D) Karyotyping analysis of both normal iPSCs and TAD-iPSCs. (E) in vitro EB-mediated differentiation of iPSCs into three germ layers, shown by immunofluorescence staining of AFP (Endoderm), α-SMA (Mesoderm), TUJ1 (Ectoderm). Scale bar=200um. (F) Hematoxylin and eosin (HE) staining of teratoma derived from both normal and TAD iPSCs, Scale bar=50um.

### 3.2 SMC differentiation from TAD-iPSCs

VSMCs play a central role in the pathogenesis of TAD^12^. To investigate the functions of VSMC in TAD, we differentiated both TAD-and normal-iPSCs into VSMCs following our recent protocol^22^. Initially, iPSC lines were differentiated into mesodermal cells over three days and subsequently induced into VSMCs using PDGF-BB and activin A in another three days (Fig.2A). Throughout this process, we did not observe a significant difference in the differentiation potential between the two groups. Both normal-and TAD-iPSC lines showed increased expression of the SMC marker genes, including α-SMA, SM22α, and calponin (CNN), along with decreased expression of pluripotency marker OCT4, as determined by RT-qPCR (Fig.2B). We further verified the increased expression of VSMCs genes α-SMA, SM22α, and CNN through immunoblotting (Fig.2C). The differentiated VSMCs were maintained in SMGM, and the expression of VSMC differentiation markers was further confirmed by immunofluorescence staining (Fig.2D). Additionally, flow cytometry analysis showed that the majority of differentiated cells expressed α-SMA (Fig.2E).

**Fig. 2.**
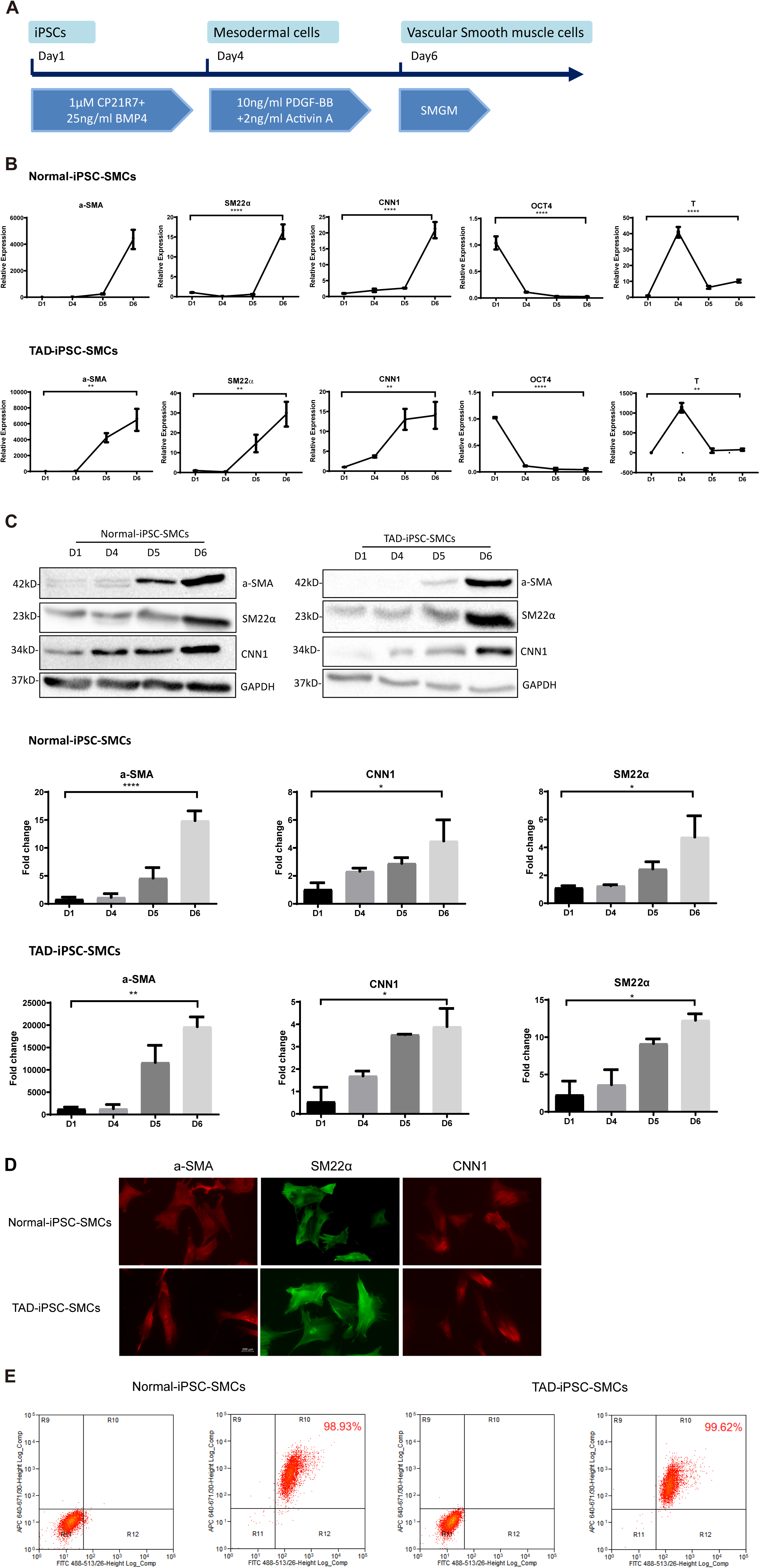
Mesoderm-mediated SMC differentiation. (A) Schematic illustration of the SMC differentiation from iPSCs. (B) SMC marker gene expression during SMC differentiation from normal and TAD-iPSCs. (C) Protein expression of SMC marker genes during SMC differentiation from normal and TAD-iPSCs by West blotting. (D) immunofluorescence staining of SMC marker genes in normal and TAD-iPSCs. (E) Flow cytometry results for iPSC-SMC differentiation. Data are shown as the mean ± S.D. for at least three independent experiments. *P<0.05, ** P <0.01.

### 3.3 TAD-iPSC-SMCs reduced the contractile ability and increased proliferation rate

Previous studies have reported that SMC from TAD patients exhibit dysfunction^23^. To investigate whether TAD-iPSC-derived SMCs (TAD-iPSC-SMCs) exhibit similar issues, we first examine their contraction ability. of TAD-iPSC-SMCs is dysfunctional. The cells were treated with carbachol, an agonist against acetylcholine receptors, for 30min. This treatment induced contraction, which resulted in a change in cell surface area on the dishes. TAD-iPSC-SMCs showed a lower contractile rate in response to carbachol compared to normal-iPSC-SMCs (Fig. 3A and 3B). Consistently, both the protein and mRNA expression levels of SMC contractile proteins, including α-SMA, SM22α, and CNN, were decreased in TAD-iPSC-SMCs compared to normal-iPSC-SMCs (Fig. 3C and 3D).

**Fig. 3.**
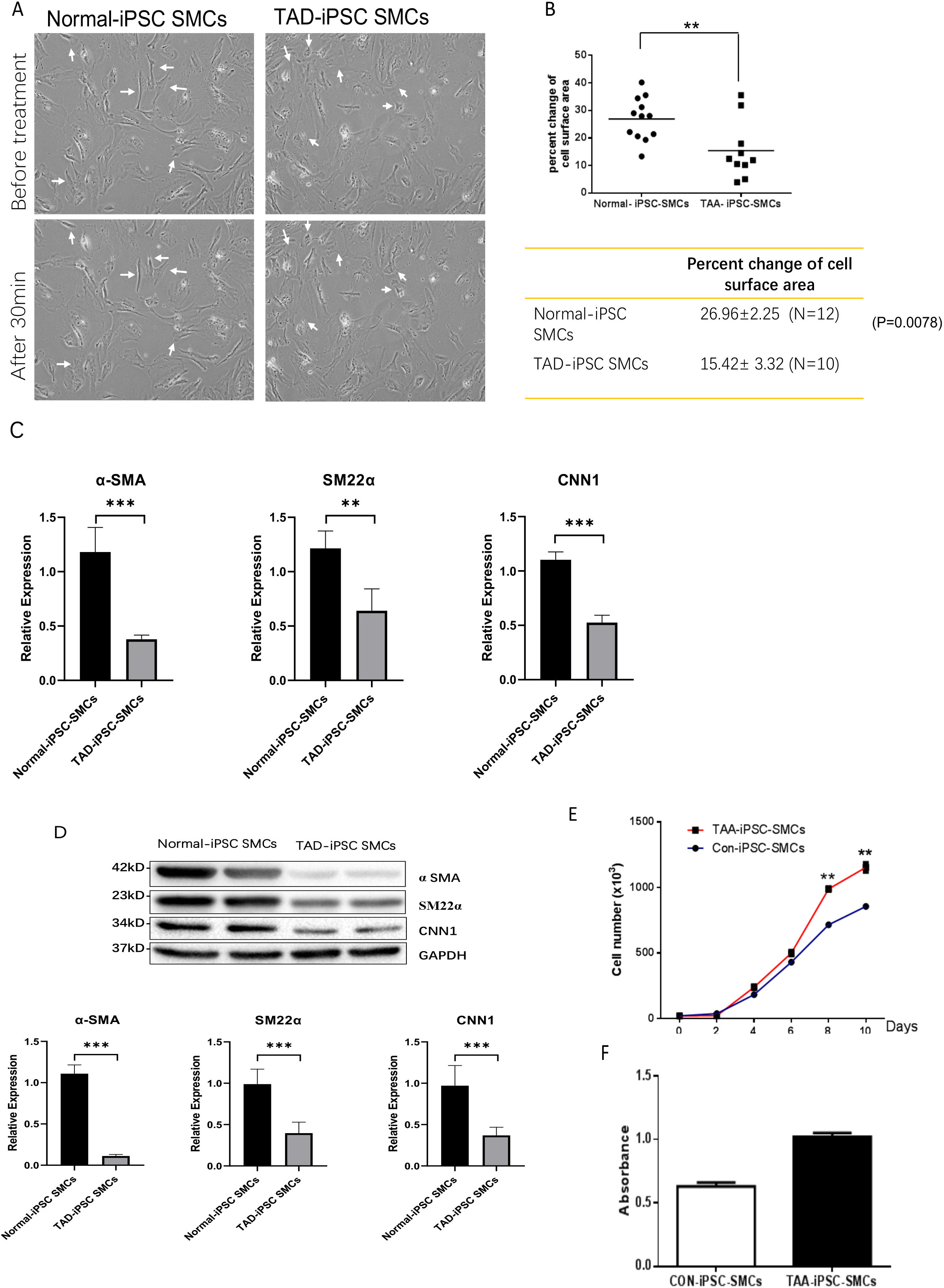
TAD-iPSC-SMCs reduce contractile ability and increase proliferation compared to control-iPSCs-SMCs. (A) TAD-iPSC-SMCs reduce contractile ability compared to normal-iPSCs-SMCs. TAD-iPSC-SMCs were treated with carbachol (1mM) and images were recorded every 1 min for 30 consecutive minutes. (B) Quantitate iPSCs-derived SMC contraction assay. (C) Comparison of SMC marker gene expression levels in normal and TAD-iPSC-SMCs. (D) Immunoblot analysis and quantification of SMC marker gene expression in normal and TAD-iPSC-SMCs. (E) TAD-iPSC-SMCs show a higher proliferation rate compared to normal-iPSC-SMCs, as indicated by the growth curve. (F) MTT assay confirmed the higher proliferation ability in TAD-iPSC-SMCs. Data are shown as the mean ± S.D. for at least three independent experiments. *P<0.05, ** P <0.01.

Additionally, TAD-iPSC-SMCs exhibited a greater proliferation capacity than normal-iPSC-SMCs, as demonstrated by the proliferation curve and confirmed by MTT assay (Fig. 3E and 3F). These results suggest that SMCs in TAD-iPSC-SMCs may undergo phenotypic switching, consistent with previous reports ^24^.

### 3.4 TAD-iPSC-SMCs show abnormal collagen expression

To identify the potential genetic defect underlying the abnormal function of TAD-iPSC-SMCs, we performed whole exome sequencing of PBMCs from the TAD patient. Intriguingly, we found a mutation in the COL4A2 gene among several predicted impaired mutations that we suspected might be associated with TAD. Sanger sequencing confirmed the COL4A2 R131M (c.392 G>T) mutation in both TAD-iPSCs and the derived SMCs (Fig. 4A). The R131 residue of COL4A2 is relatively conserved across several mammalian species (Fig. 4B). COL4A2 encodes the collagen type IV α2 chain, and its mutations have been linked to various cardiovascular diseases, including aneurysms^25, 26^. Using a web-based prediction tool, we found that the COL4A2 R131M mutation is predicted to be harmful (Fig. 4B). We then examined the expression of several vascular collagen genes in both normal-and TAD-iPSC-SMCs. Our results showed that the expression of COL4A2 and COL4A1 was decreased in TAD-iPSC-SMCs compared to normal iPSC-SMCs, while the expression of COL1A1 and COL3A1 genes was increased (Fig. 4C-D). When SMCs were exposed to TGF-β, all collagen genes except for COL4A2 showed increased expression (Fig. 4C). These findings suggest that the COL4A2 mutation might contribute to the phenotype observed in TAD-iPSC-SMCs.

**Fig. 4.**
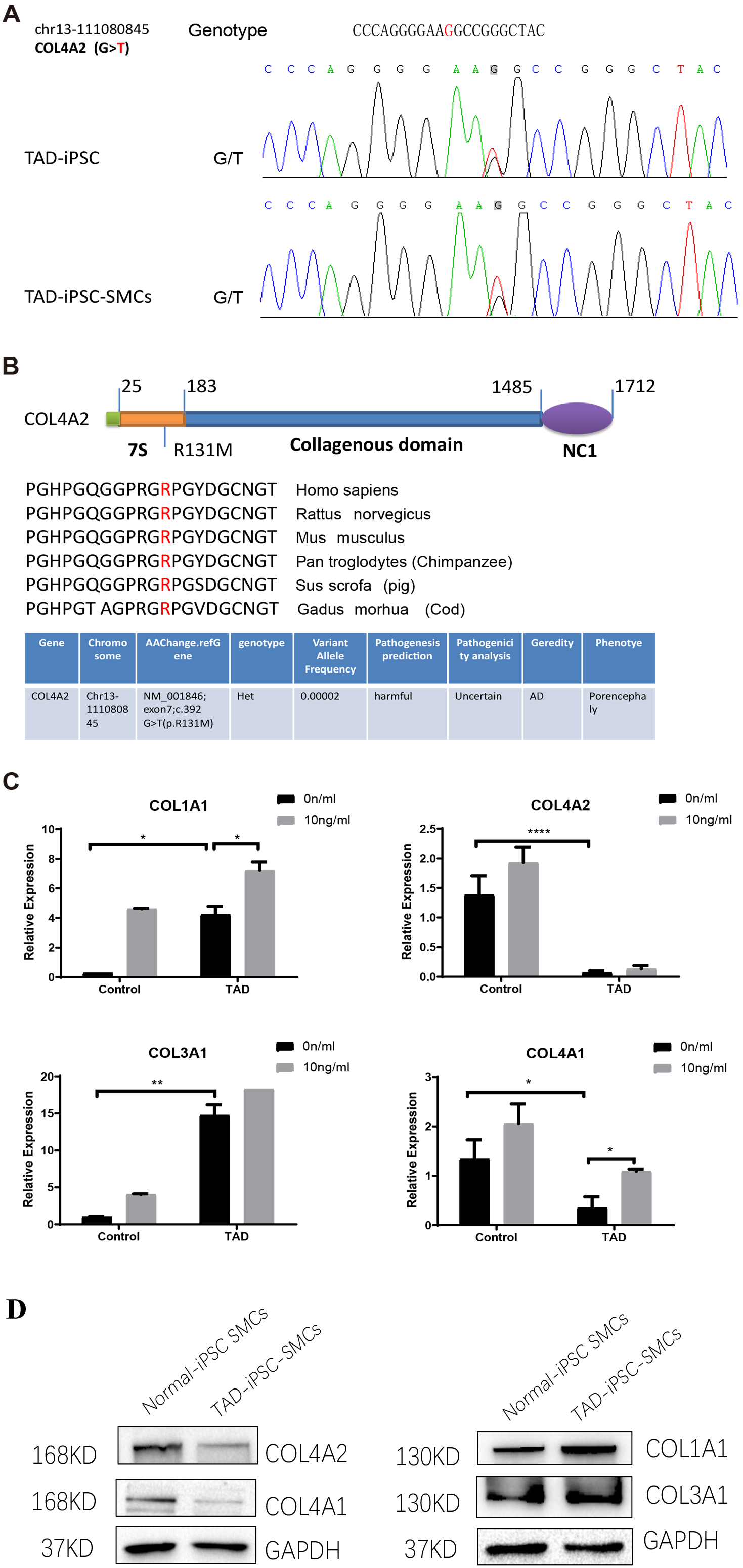
TAD-iPSC and derived SMCs carry a mutation in collagen type IV (*COL4A2)*. (A) Sanger sequencing of collagen IV A2 (COL4A2) mutation (c.392G>T, p. R131M) in the TAD-iPSC line and the derived SMCs. (B) Schematic structure of COL4A2 showing the mutation R131M located in the 7S domain, which is relatively conserved among several mammal species. (C) Expression of various types of collagen in both normal and TAD-iPSC-SMCs, with and without TGF-β treatment. (D) Protein expression level of COL4A2 and COL4A1 decreases, while COL1A1 and COL3A1 increased. Data are shown as the mean ± S.D. for at least three independent experiments. *P<0.05, ** P <0.01.

### 3.5 Noncanonical TGF-**β** signaling activated in TAD-iPSC-SMCs

To determine whether TGF-β signaling contributes to the progression of the TAD phenotype, we examined both the canonical TGF-β-Smad and noncanonical TGF-β-ERK signaling pathways in TAD-iPSC-SMCs. We observed that the level of phosphorylated ERK1/2 (pERK1/2), but not phosphorylated SMAD2 (pSMAD2), was higher at the basal level in TAD-iPSC-SMCs compared to normal iPSC-SMCs (Fig. 5A). Additionally, TAD-iPSC-SMCs exhibited less pSMAD2 activation following TGF-β stimulation.

**Fig. 5.**
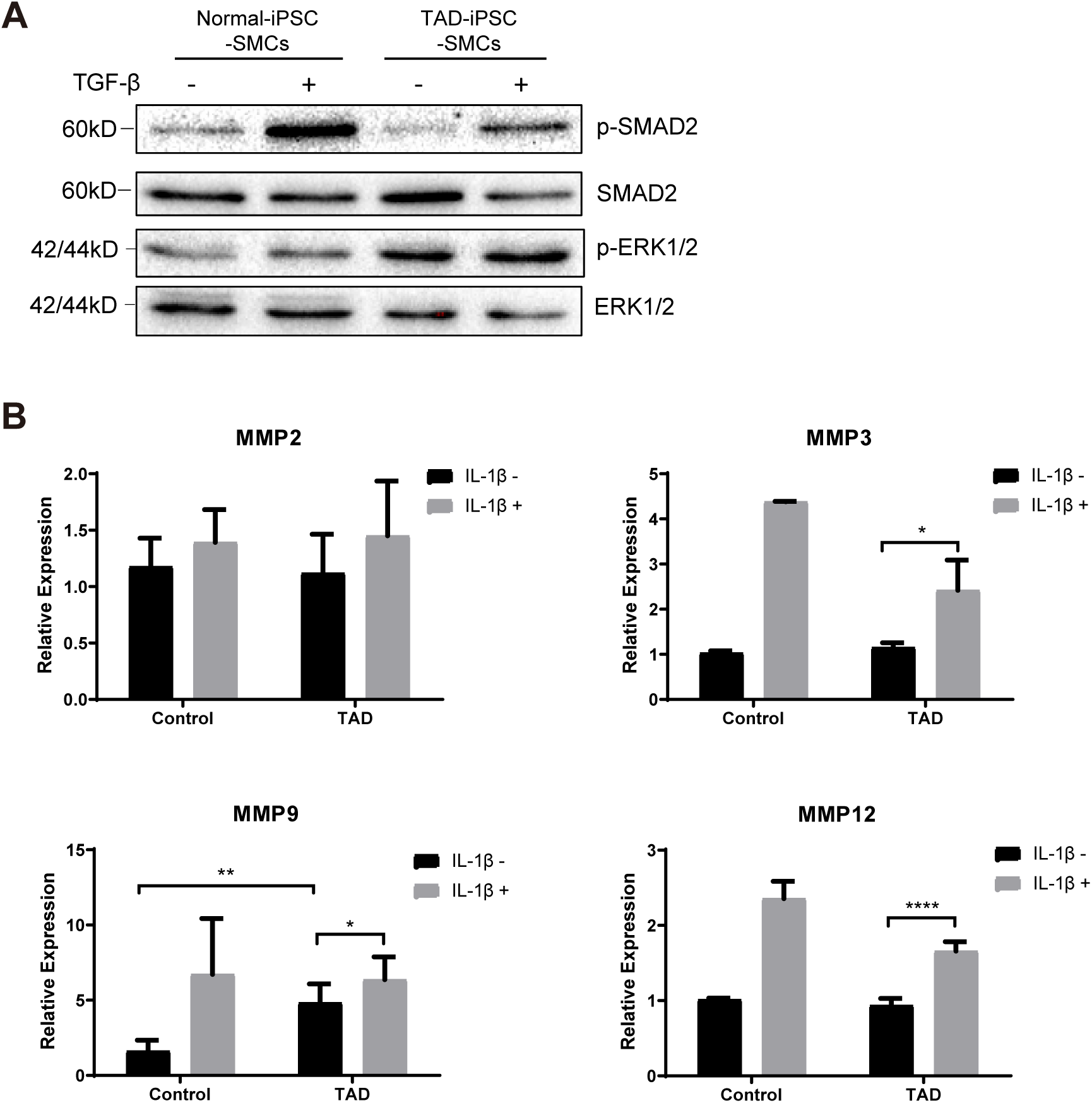
Noncanonical TGF-β signaling is activated and MMP9 expression is increased in TAD-iPSC-SMCs. (A) Western blotting analysis of phosphorylated TGF-β canonical (SMAD2) and noncanonical (ERK1/2) pathways in normal and TAD-iPSC-SMCs with or without TGF-β treatment. (B) RT-qPCR expression of MMPs in normal and TAD-iPSC-SMCs with or without IL-1β (10ng/ml) treatment. Data are shown as the mean ± S.D. for at least three independent experiments. *P< 0.05, ** P <0.01.

We also compared the expression levels of matrix metalloproteinases (MMPs) in normal- and TAD-iPSC-SMCs. Among the MMPs examined, only MMP9 showed relatively higher expression in TAD-iPSC-SMCs. However, treatment with the cytokine IL-1β for 24 hours elevated the mRNA expression levels of MMP2, MMP3, MMP9, and MMP12 in both normal and TAD-iPSC-SMCs (Fig. 5B). These results suggest that TAD-iPSC-SMCs exhibit an imbalance between the canonical Smad2-dependent and noncanonical ERK1/2-dependent TGF-β signaling pathways.

## 4 Discussion

In this study, we established iPSCs from a TAD patient and differentiated them into SMCs to create a disease cellular model for investigating the molecular mechanisms of human TAD. Using this patient-specific iPSC and SMC differentiation platform, we discovered that TAD-iPSC-SMCs exhibited reduced contraction ability and increased proliferation rate, indicative of phenotypic switching as early events in aortic dissection. Additionally, we identified a COL4A2 mutation in this patient, with TAD-iPSC-derived SMCs showing reduced COL4A2 expression but increased expression of COL1A1 and COL3A1. Furthermore, our model revealed dysfunctional TGF-β signaling, characterized by inhibited canonical Smad2-mediated pathways and activated noncanonical ERK1/2-mediated pathways in SMCs.

The derivation of human iPSCs has provided unprecedented opportunities for cardiovascular disease modeling and regenerative medicine^27^. Previous studies have demonstrated the application of iPSCs in modeling various cardiovascular diseases, such as long-QT syndrome and arrhythmogenic right-ventricular dysplasia ^28, 29^. Recently, the Sinha group developed an in vitro Marfan syndrome (MFS)-iPSC model, using iPSC-derived smooth muscle cells to elucidate the role of TGF-β signaling in MFS pathogenesis, and identified a novel role of p38 and KLF4 in the MFS pathogenesis ^18^. Our study extends this approach to thoracic aortic dissection, demonstrating that TAD-iPSC-SMCs faithfully mimic human SMC phenotypes associated with TAD. This model adds to the growing list of iPSC-based disease models and offers a potential platform for screening novel therapeutics for TAD.

The extracellular matrix (ECM), particularly collagen, is crucial in the pathogenesis of TAD. Collagen I and III, the most abundant fibrous collagens in the aortic wall, are essential for tensile strength and cytokine sequestration. Dysfunction of collagen III is linked to TAD in Ehlers-Danlos syndrome (vascular type IV, OMIM#130050). Although some studies have reported increased collagen I and III expression in TAD, de Figueiredo Borges et al. observed decreased collagen in the outer half of the dissected aortic wall ^30^, suggesting that collagen content in the media may decrease prior to dissection. Interestingly, type IV collagen, a basement membrane-specific type, is also implicated in TAD. Microarray data from the Sandmann group identified downregulation of COL4A2 and COL4A5 in aortic dissection^11^ .

Consistent with these findings, we observed decreased COL4A2 expression in TAD-iPSC-SMCs and identified a COL4A2 R131M mutation in the patient. This mutation may weaken the tunica intima by reducing collagen IV expression, contributing to TAD formation. Variants of collagen genes, including COL4A2, have been implicated in sporadic AD pathogenesis^31^ , and COL4A1 and COL4A2 mutations are associated with a broader spectrum of cardiovascular, renal, ophthalmological, and muscular abnormalities as well as aneurysms ^26, 32^. A recent study reported that a loss-of-function mutation in lysyl oxidase (LOX), an enzyme critical for collagen cross-linking, causes thoracic aortic aneurysm and dissection^33^, supporting our hypothesis that mutations in collagen-encoding genes may weaken the aortic wall and lead to dissection formation.

In healthy vascular walls, SMCs maintain vascular tone and remain quiescent in the contractile phenotype. Under pathological conditions, SMCs can dedifferentiate into a synthetic phenotype and proliferate. This phenotypic switching is recognized as an early event in thoracic aortic aneurysm formation^14^, characterized by downregulation of VSMC differentiation markers such as SM22α and smooth muscle α-actin . Our findings in TAD patient-specific iPSC-SMCs are consistent with this observation (Fig.3C-D). This is consistent with our current result in the TAD patient-specific iPSC-SMC. The Milewicz group reported that TGFBR2 mutations lead to decreased expression of SMC contractile proteins, suggesting that defective SMC contractile function contributes to TAAD pathogenesis ^34^. Although the underlying mechanisms remain unclear, recent research indicates the pivotal role of the XBP1u-FoxO4-myocardin axis in maintaining the VSMC contractile phenotype and preventing phenotypic switching during aortic aneurysm formation ^35^. ER stress pathways, potentially involved in aortic dissection or aneurysm, may also be implicated, as suggested by previous studies^36^ . Although COL4A2 mutations may also cause ER stress, we did not investigate this pathway in our study.

TGF-β signaling plays a crucial role in TAD development. This multifunctional signaling molecule is involved in various physiological processes, including SMC differentiation, ECM synthesis, and degradation via upregulation of MMPs. TGF-β activates both Smad2/3-mediated canonical pathways and non-Smad-mediated (noncanonical) pathways. Initial studies suggested that excessive TGF-β signaling contributes to thoracic aortic dissection and aneurysm ^37^. Habashi et al. reported that the onset of thoracic aortic disease in Fbn1C1039/+ mice is associated with increased TGF-β signaling, which can be prevented by TGF-β inhibition or AT1R blockade with losartan ^38^. Patients with mutations in genes such as α-SMA, MYH11, and fibulin-4 also exhibit TGF-β hyperactivity ^39^. Furthermore, patients and mice with heterozygous loss-of-function mutations in genes encoding components of canonical TGF-β signaling pathway, such as ligands, receptors, and signal transducers also found overactivation of TGF-β signaling. Conversely, recent evidence suggests that TGF-β signaling may protect against aortic aneurysm and dissection. Li et al. and Wei et al. found that Tgfbr2 disruption in smooth muscle cells exacerbates aneurysm and dissection in Marfan syndrome mice , indicating that TGF-β signaling is essential for postnatal aortic growth and homeostasis ^40, 41^. TGF-β could activate intracellular smad-dependent canonical pathway and smad-independent noncanonical pathways (e.g. ERK1/2, JNK). While inhibition of noncanonical TGF-β signaling ameliorates aortic aneurysm progression in Marfan syndrome mice^42^, our iPSC-SMC model showed hyperactivation of noncanonical ERK1/2 signaling and reduced Smad2 activity in response to TGF-β in patient-derived SMCs (Fig. 5A). (Fig.5A). This imbalance between noncanonical and canonical TGF-β signaling likely contributes to aortic dissection or aneurysm development.

However, our study has some limitations. For example, further data are needed, such as applying genome editing techniques to correct the COL4A2 mutation and confirming the reversal of SMC dysfunction after mutation correction.

In summary, our work has modeled the TAD-associated SMC mechanism using patient-specific iPSCs. Our study revealed that TAD-iPSC-SMCs with a COL4A2 mutation exhibit phenotypic switching and activation of noncanonical ERK1/2 TGF-β signaling, contributing to TAD progression. Our findings highlight the potential importance of the noncanonical TGF-β pathway in TAD pathogenesis and suggest that targeting this pathway could be a therapeutic strategy for TAD treatment in the future.

## Ethics statement

This study was approved by the medical research ethics committee of Wenzhou Medical University (approval number: 2017-066). The study was conducted in accordance with the local legislation and institutional requirements.

## Conflict of Interest

The authors declare that the research was conducted in the absence of any commercial or financial relationships that could be construed as a potential conflict of interest.

## Author contributions

PJ: Formal Analysis, Investigation, Methodology, Visualization, Writing–original draft, Writing–review and editing. YX: Formal Analysis, Data Curation, Investigation, Writing– original draft, Writing–review and editing. SW: Formal Analysis, Data Curation, Investigation, Writing–review and editing. LD: Data curation, Formal Analysis, Writing–review and editing. YC: Investigation, Writing–review and editing. MZ: Investigation, Writing–review and editing. XC: Funding acquisition, Investigation, Methodology, Writing–review and editing. XF: Project administration, Resources, Writing–review and editing. YG: Supervision, Validation, Writing– review and editing. ML: Supervision, Validation, Writing–review and editing. YW: Conceptualization, Visualization, Funding acquisition, Investigation, Supervision, Writing– original draft, Writing–review and editing.

## Funding

The author(s) declare financial support was received for the research, authorship, and/or publication of this article. This work was supported by the National Natural Science Foundation of China (82070487, 32370787,81670454), and the Scientific Research Start-up Fund of Wenzhou Medical University (QTJ15029), Wenzhou Major Science and Technology Innovation Project (ZY2021014).

## Acknowledgments

We would like to thank all the members of the Institute of Hypoxia Medicine, the patient, and his family who contributed to this work. We appreciate the great help from Dr. Jiaxi Zhou (State Key Laboratory of Experimental Haematology in China) for this project.

---

**Fig.S1**

The iPSC reprogramming process with morphology changes at different stages from primary VSMC to established iPSC colonies. Scale bar=250μm.

**Fig.S2**

The cell morphology changes during mesoderm-mediated SMC differentiation from iPSCs. Scale bar=250μm.

## Notes

### Competing Interest Statement

The authors have declared no competing interest.

