## Supplementary figures and images for "Modeling Thoracic Aortic Dissection Using Patient-Specific iPSCs Reveals VSMC Dysfunction and Extracellular Matrix Dysregulation"

### 02 Figs supplementary 2024-10-06.pdf

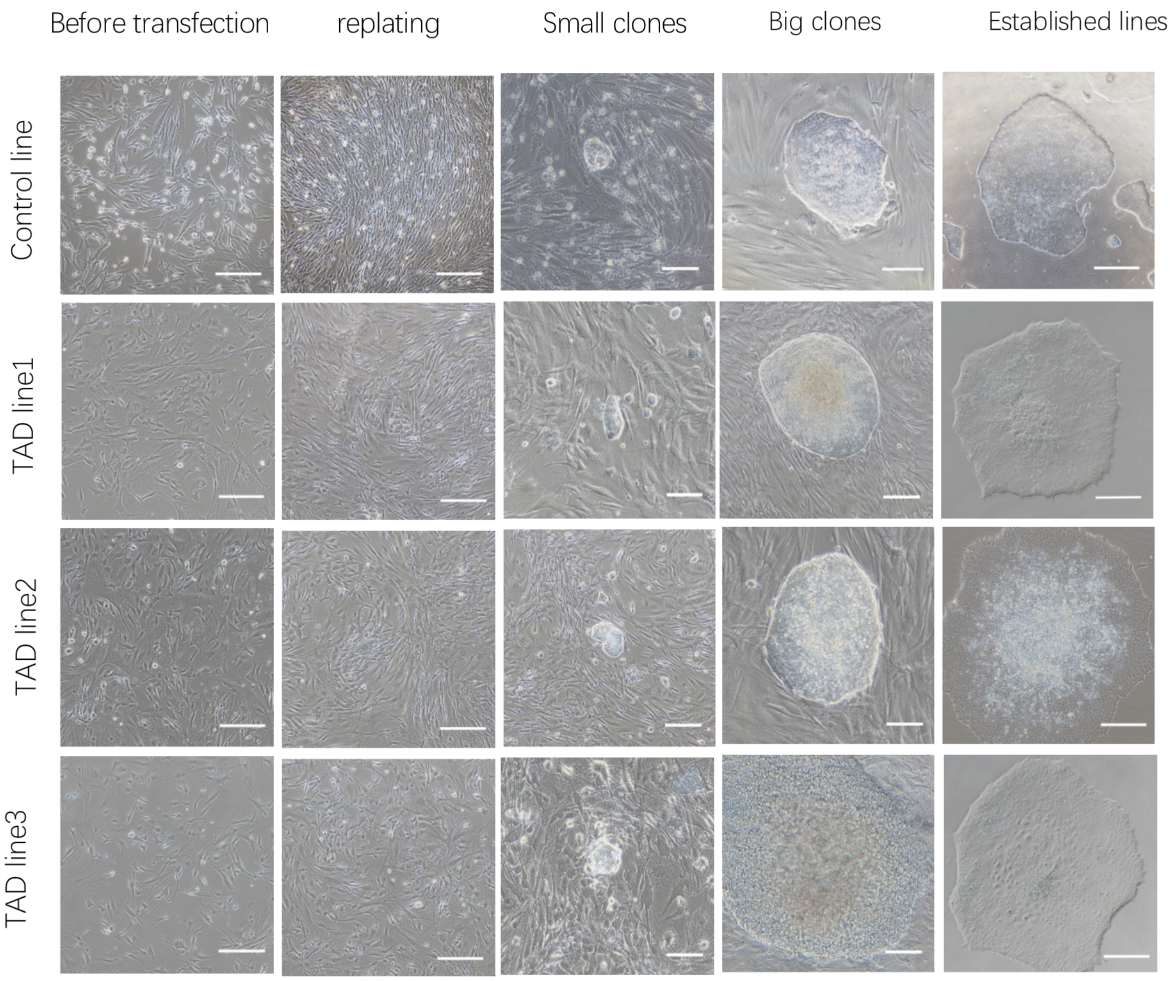

Fig S2

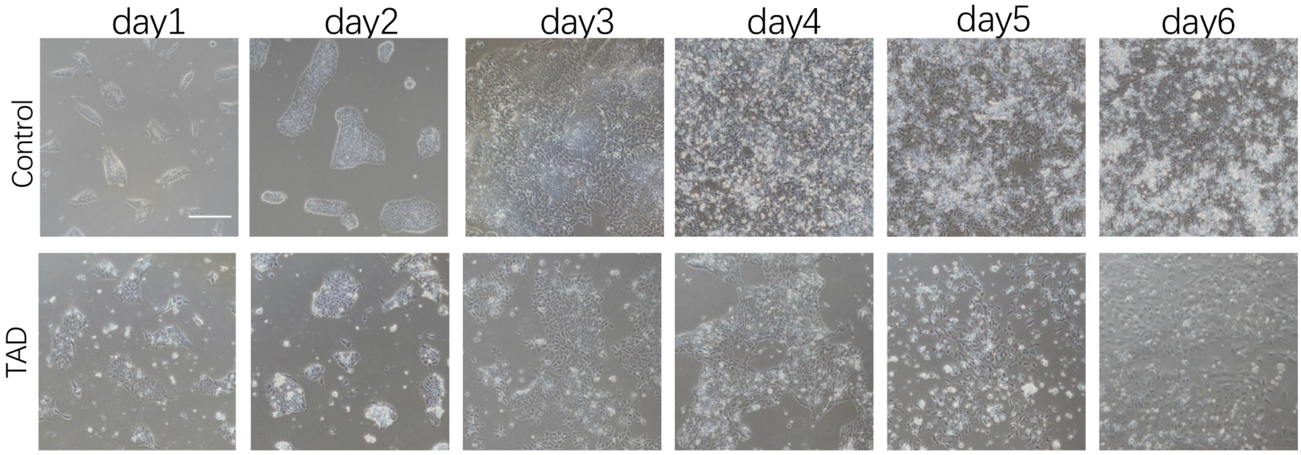


Bar=100um
